# Mnemonic content and hippocampal patterns shape judgments of time

**DOI:** 10.1101/2021.08.03.454949

**Authors:** Brynn E. Sherman, Sarah DuBrow, Jonathan Winawer, Lila Davachi

**Affiliations:** Department of Psychology, Yale University; Department of Psychology, University of Oregon; Department of Psychology and Center for Neural Science, New York University; Department of Psychology, Columbia University; Nathan Kline Institute for Psychiatric Research

## Abstract

Our experience of time can feel dilated or compressed, rather than reflecting true “clock time.” Although many contextual factors influence the subjective perception of time, it is unclear how memory accessibility plays a role in constructing our experience of and memory for time. Here, we used a combination of behavioral and fMRI measures to ask the question of how memory is incorporated into temporal duration judgments. Behaviorally, we found that event boundaries, which have been shown to disrupt ongoing memory integration processes, result in the temporal compression of duration judgments. Additionally, using a multivoxel pattern similarity analysis of fMRI data, we found that greater temporal pattern change in the left hippocampus within individual trials was associated with longer duration judgments. Together, these data suggest that mnemonic processes play a role in constructing representations of time.

**Statement of Relevance:** Our everyday experiences convey a powerful truth: That our *perception* of time often diverges from the *reality* of time. When enjoying an active vacation with family, time moves quickly: hours go by in minutes. When sitting through an unnecessary meeting, time moves slowly: minutes go by in hours. What is the origin of these phenomenologically compelling illusions of time perception? Past research has examined how a range of specific factors, from emotions to blinking, contribute to the distortion of time. Here, in contrast, we evaluate how the content and accessibility of our memories shapes time perception. We show that context shifts, known to disrupt memory processing, also lead to robust contractions of perceived time. We discuss how both effects — memory disruptions and time distortions — may be linked via the hippocampus.

“An hour, once it lodges in the queer element of the human spirit, may be stretched to fifty or a hundred times its clock length; on the other hand, an hour may be accurately represented on the timepiece of the mind by one second.” — Virginia Woolf

## Introduction

On a busy vacation, time may escape you — by the time you go to the museum, grab lunch in the park, souvenir shop, and visit a historical site, the day may seem to have flown by. Yet, when recalling the trip to a friend, that same day may feel like a week; all of those events could not have possibly occurred within the same few hours. This puzzle raises a fundamental question of how the structure of experience can paradoxically influence subjective impressions of time in experience and in reflection.

A great deal of work has focused on the latter: how the structure of experience influences how we *remember* elapsed time. In particular, abrupt shifts in context, or event boundaries, influence memory for time. Memory for the temporal order of events is disrupted across event boundaries (DuBrow & Davachi, 2013; Horner et al., 2016; Heusser et al., 2018). Further, intervals which contain an event boundary are remembered as *longer* than equivalently-timed intervals without a boundary, (Ezzyat & Davachi, 2014; Clewett et al., 2020) and mnemonic duration judgments scale with the number of events (Faber & Gennari, 2015; Lositsky et al., 2016; Faber & Gennari, 2017). Such findings converge with the intuition that busy days feel long in memory: events may serve to dilate time in memory. How, though, do events also result in time feeling subjectively shorter *in the moment*?

Event boundaries also influence memory on the more immediate time-scale of working memory by reducing access to information prior to the boundary (Zwaan, 1996; Rinck & Bower, 2000; Speer & Zacks, 2005; Ezzyat & Davachi, 2011). Thus, perhaps this decreased accessibility to mnemonic information translates into a *contraction* of time for intervals containing event boundaries. Models of time perception posit an “integrator” or “accumulator” which sums across experience to determine how much time has passed (e.g., Droit-Volet & Meck, 2007; Wittmann, 2013). By reducing the contents of working memory, event boundaries may reduce the information integrated into duration judgments, leading to an underestimation of time. Consistently, there is some evidence that in-the-moment time judgments are compressed after one or more discrete boundaries (Liverence & Scholl, 2012; Bangert et al., 2019; Yousif & Scholl, 2019). However, whether this effect of event boundaries on more immediate time judgments is indeed due to a decreased accessibility to the prior event remains an open question.

A separate body of literature has examined the interplay of time and memory by asking whether the hippocampus — a structure critical for episodic memory (Scoville & Milner, 1957) — plays a role in tracking time. In one seminal study, Meck and colleagues found that although disrupting hippocampal function did not result in impairments in perceiving duration, it critically impaired temporal working memory and led to underestimation of the “reference memories” (Meck et al., 1984). This study suggests that the hippocampus may be involved in duration processing, but leaves open the question of how, precisely, mnemonic information is incorporated into representations of time.

A plethora of subsequent work has established that the medial temporal lobe — in particular the hippocampus — indeed is sensitive to temporal duration information (Eichenbaum, 2013; Davachi & DuBrow, 2015; Lee et al., 2020). Independent of memory demands, populations of hippocampal neurons reflect the passage of time (Manns et al., 2007; Mankin et al., 2012; Rubin et al., 2017). Individual hippocampal neurons are sensitive to temporal information, firing during delays (MacDonald et al., 2011; 2013; Naya & Suzuki, 2011; Sakon et al., 2014; Umbach et al., 2020; Reddy et al., 2021) or at specific temporal moments (Terada et al., 2017; Sun et al., 2020). Further, memories acquired close in time are coded by overlapping populations of neurons (Cai et al., 2016). Such overlapping representations may have consequences for the subjective representation of time in *long-term memory:* events remembered as further apart in time are associated with greater fMRI pattern change in the medial temporal lobe (Ezzyat & Davachi, 2014; Nielson et al., 2015; Deuker et al., 2016; Losistsky et al., 2016). However, whether and how such hippocampal representations influence more immediate judgments of time remains unclear.

In the present study we examine the role of memory accessibility and hippocampal representations on subjective judgments of time. Critically, we induce context shifts by inserting event boundaries, allowing us to assess the role of memory representations while holding true duration constant. First, in three behavioral experiments, we extend prior work demonstrating that event boundaries reduce estimates of duration, and provide evidence that these reductions are due to decreased mnemonic accessibility. In a final fMRI experiment, we use pattern similarity to track changes in representations within a single temporal interval, and find that pattern change in the left hippocampus supports duration judgments, suggesting that the hippocampus may carry behaviorally-relevant temporal information on the order of seconds.

### Experiment 1

To examine how event boundaries affect subjective duration judgments, we designed a paradigm in which participants viewed a colored square (0.5 - 5s in Experiments 1-3; 2 - 8s in Experiment 4; **Figure 1A**), which remained a single color for the duration of the trial (continuous condition) or switched colors during the interval (boundary condition). Participants then judged how long the square, regardless of color, was presented on the screen using a continuous timeline (**Figure 1B**). We hypothesized that the color switches act as event boundaries, thus decreasing accessibility of the pre-boundary interval in memory. Critically, if memory for information from across the entire interval is integrated into a duration estimate, then reduced memory access for pre-boundary information would lead to shorter duration judgments in the boundary condition.

**Figure 1.**
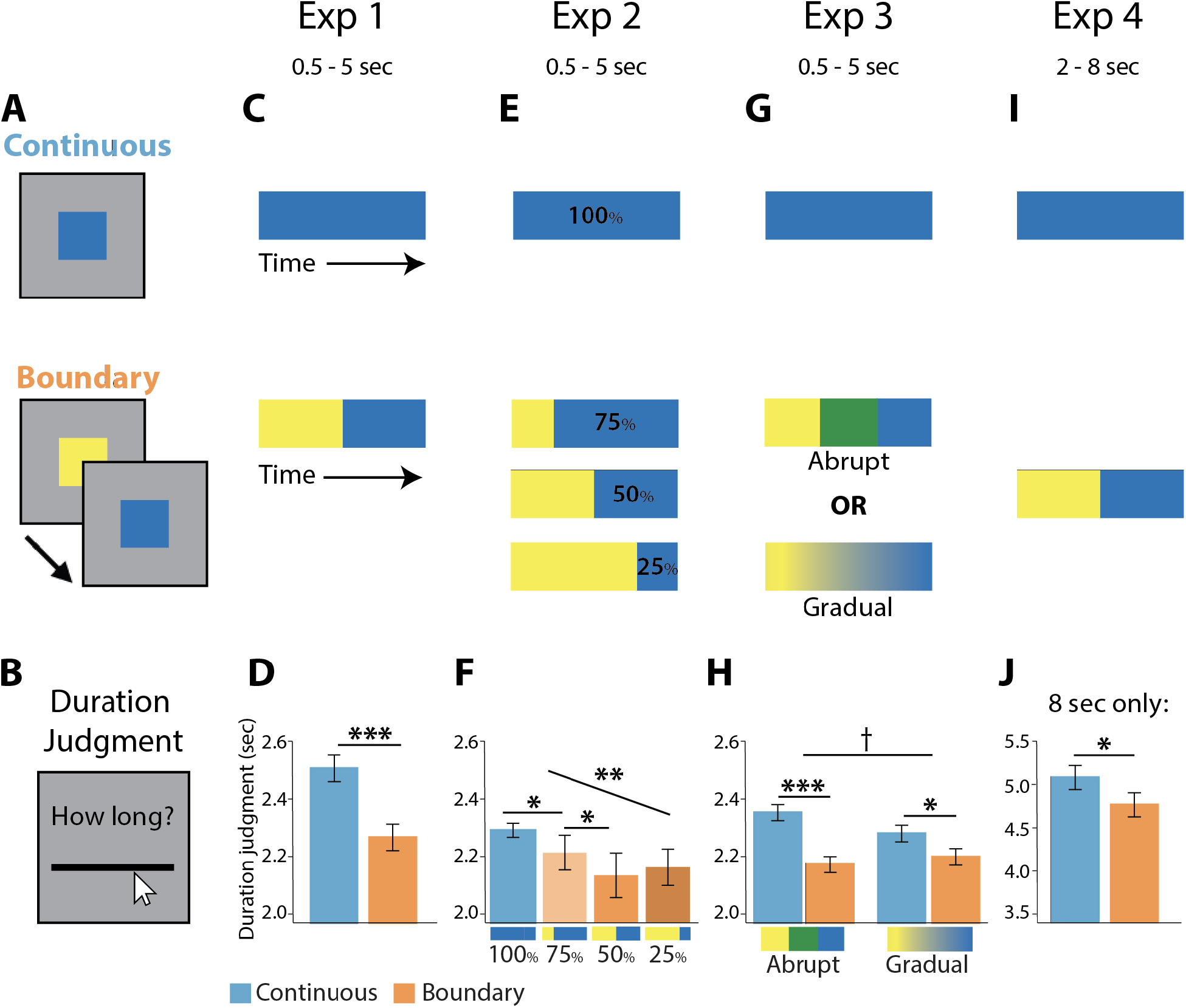
Behavioral Task & Data. A) Participants viewed a colored square, which either stayed the same color for the total duration (continuous condition) or switched colors during the interval (boundary condition). B) Participants then estimated how long the square was on the screen for (regardless of color change). C) Schematic of trial structures. In experiment 1, boundaries always occurred halfway through the total 0.5-5sec interval. D) Experiment 1 results: boundary trials were judged as significantly shorter than continuous trials. E) In experiment 2, color switches in boundary conditions occurred either one-quarter, one-half, or three-quarters through the total 0.5-5sec interval, such that the second event was 75%, 50%, or 25% of the total duration, respectively. F) Experiment 2 results: there was a significant linear effect of boundary placement; pairwise comparisons showed the 75% trials were judged to be reliably shorter than 100% trials, and 50% trials were judged as reliably shorter than 75% trials. G) In experiment 3, boundary trials were either abrupt (two color switches, each color presented for one-third of the 0.5-5sec interval) or gradual (smoothly morphing through color space for the entire 0.5-5sec interval). H) Experiment 3 results: in both groups, boundary trials were judged as reliably shorter than continuous trials, though this effect was weaker in the gradual change group. I) As in experiment 1, boundaries occurred halfway through the interval, though the interval was extended to be 2-8 sec. J) Experiment 4 results: boundary trials were judged as reliably shorter than continuous trials. *** p < .001, two-tailed; ** p < .005, two-tailed; *p < .05, two-tailed; † p < .07, two-tailed. Error bars denote the within-participant standard error of the mean.

## Methods

### Participants

Twenty-one individuals (13 female; age range 18-33; mean = 21.5) were recruited from New York University and the larger community and participated for either course credit or payment ($10/hr). Informed consent was obtained in a manner approved by the University Committee on Activities Involving Human Subjects at New York University. We aimed to collect data from twenty usable participants; one participant was excluded for reporting that they explicitly counted the intervals during the task, and thus one additional participant was collected.

### Stimuli

Stimuli were colored blue (R = 0, G = 0, B = 255), green (R = 0, G = 255, B = 0), and yellow (R = 260, G = 200, B = 80) squares presented centrally on a mid-gray background. Each square was presented on-screen for a 0.5 – 5 seconds interval (sampled equally in increments of 0.5 seconds). For half of the trials, the color remained the same for the entire duration (continuous condition), and for the other half of the trials the color switched half-way through the total duration (boundary condition).

Each participant viewed each color/color-pair once for each time point. For each participant, color pairings were randomly assigned such that the pairing of the three colors in the boundary condition was fixed (for example, the boundary trials for one participant would always be blue-green, green-yellow, yellow-blue, and this pairing would be randomly assigned for each participant). The order of presentation was pseudo-randomized such that condition-duration combinations were not repeated back-to-back and no condition or duration appeared more than 4 times consecutively.

In total, the task consisted of 3 trials per condition and time point (60 trials total).

### Duration Judgment Task

On each trial, participants were instructed to attend to the square and keep track of the time. Importantly, participants were explicitly verbally instructed not to count while the square was on the screen (Rattat & Droit-Volet, 2012), and debriefing questionnaires suggested that compliance with this instruction was high.

Participants subsequently were presented with a continuous timeline with the prompt “How long was the square on the screen?” This timeline was bounded by 0.5 sec and 5 sec. Participants were instructed to disregard any color changes and estimate the total duration of the square presentation, regardless of color. Participants responded using a computer mouse. Responses were self-paced.

## Results

To assess accuracy of participants’ duration judgments, we calculated a Spearman rank correlation between the actual duration of the trial and the participant’s subjective judgment (response made on the number line) for each participant. For the purposes of statistical evaluation, rho values were Fisher-transformed. All participants exhibited a positive Spearman rank correlation value; Mean correlation, or z = 1.29, 95% CI = [1.15, 1.43], *t*(19) = 19.14, *p* < 0.001, *d =* 4.28 (**Supplemental Figure 1A)**.

To further confirm that each participant performed above chance relative to their own response distribution (i.e., irrespective of response bias), a permutation test was also computed for each participant: over 10,000 iterations, each participant’s full distribution of responses was shuffled, and the rank correlation between the time of each trial and this shuffled response was computed. Significance was determined as being higher than 95% of this null distribution. All participants in all experiments demonstrated accuracy reliably above chance relative to their own distributions.

Next, we addressed our key question that event boundaries influence temporal duration judgments. In this experiment, in which boundary trials switched colors half-way through (**Figure 1C**), we found that participants judged boundary trials to be reliably shorter than continuous trials (Mean difference, M = −0.24, 95% CI = [ −0.34, −0.14]), *t*(19) = −5.07, *p* < 0.001, *d* = −1.13. (**Figure 1D**). This finding conceptually replicates prior findings demonstrating a compression of duration judgments for experiences which contain a boundary (Liverence & Scholl, 2012; Bangert et al., 2019; Yousif & Scholl, 2019).

### Experiment 2

After establishing an influence of event boundaries on duration judgments in this task, we next aimed to understand *why* boundaries lead to the compression of time. One reason why boundary trials might be judged as shorter is because the pre-boundary color is less accessible in working memory (Zwaan, 1996), leaving mostly the post-boundary memory to be integrated into the duration judgment, thus leading to a shorter estimate. If this “flushing” account is true, then judgments of time should be influenced not merely by the presence of a boundary, but should be sensitive to *when* in the interval the boundary occurred. In other words, boundaries which occur earlier in the trial will have more mnemonic content (as relatively less information was flushed at the boundary), and thus may be judged as longer than trials in which the boundary occurred later in the sequence.

## Methods

### Participants

Twenty-nine individuals (18 female; age range 18-24; mean = 19.4) were recruited from New York University for course credit. Informed consent was obtained in a manner approved by the University Committee on Activities Involving Human Subjects at New York University. The sample size for this experiment was chosen to be larger than that of Experiment 1 to increase sensitivity, given that more conditions were present in this experiment.

### Stimuli

Stimuli were colored blue (R = 0, G = 0, B = 200), green (R = 75, G = 205, B = 75), and yellow (R = 260, G = 200, B = 80) squares presented centrally on a mid-gray background. Each square was presented on-screen for a 0.5 – 5 seconds interval (sampled equally in increments of 0.5 seconds).

For half of the trials, as in Experiment 1, the color remained the same for the entire duration (continuous condition). The other half of the trials were one of three boundary conditions: for one third of the boundary trials (one-sixth of the total experiment), the color switched one-quarter of the way through the total duration (such that the second event was 75% of the total duration); for one third, the color switched one-half of the way through (second event was 50% of total duration); for one third, the color switched three-quarters of the way through the total duration (second event was 25% of total duration).

Each participant viewed each color/color-pair an equal number of times for each time point (here, twice over the entire experiment). Color pairings and pseudorandomization of trial sequences was conducted using the same procedure as in experiment 1.

In total, the task consisted of 3 trials per condition and time point (120 trials total).

### Duration Judgment Task

The duration judgment task was identical to that described in Experiment 1.

## Results

Again, we first assessed accuracy of participants’ duration judgments, and found that all participants exhibited a reliable correlation, mean z = 1.18, 95% CI = [1.08, 1.27], *t*(28) = 24.66, *p* < 0.001, *d* = 4.58 (**Supplemental Figure 1B)**.

In this experiment, we manipulated the placement of the boundary, such that the last event comprised 100% (continuous condition), 75%, 50%, or 25% of the trial (**Figure 1E**). First, we replicated the result of Experiment 1, such that boundary trials were judged on average to be shorter than continuous trials (M = −0.12, 95% CI = [-0.17, −0.07]), *t*(28) = −5.00, *p* < 0.001, *d* = −0.93. To test our primary hypothesis that the extent of compression is modulated by the duration of the second color event, we examined whether the position of the boundary significantly modulated participants’ duration judgments (See **Figure 1F**). We found a significant main effect of boundary placement (continuous (100%), 75%, 50%, 25%) on duration judgment (F(3, 84) = 7.69, p < 0.001, *η*_*p*_^*2*^ *=* 0.22). To assess whether this main effect reflected the scaling of duration judgments with the length of the post-boundary event, we asked whether duration judgments were linearly related to the length of the last event. For each participant, we computed the Pearson correlation between their duration judgments and the relative placement of the boundary, and found a reliable negative correlation across participants (Mean fisher-transformed correlation = −0.57, 95% CI = [-0.90, −0.25], *t*(28) = −3.66, *p* = 0.001, *d* = −0.68). To further probe this relationship, we performed planned pairwise comparisons between neighboring conditions. The 75% boundary trials were rated as significantly shorter than 100% (continuous) trials (M = −0.07, 95% CI = [-0.14, −0.02], *t*(28) = −2.64, *p* = 0.013, *d* = −0.49). The pairwise difference between 75% and 50% trials also reached significance, such that 50% trials were judged as significantly shorter than 75% trials (M = −0.08, 95% CI = [-0.16, −0.00], *t*(28) = - 2.09, *p* = 0.046, *d* = −0.39). However, we did not find a significant difference between 50% and 25% trials (M = 0.03, 95% CI = [-0.05, 0.11], *t*(28) = 0.77, *p* = 0.446, *d* = 0.14). These data suggest that subjective time scales with the length of the most recent event up to the halfway point. Thus, once the most accessible information becomes too short (e.g, half the length of the trial), participants may rely on additional heuristic information (e.g., that multiple events occurred).

To further probe this memory accessibility account, we assessed the relative contributions of the first and second event durations on the resulting duration judgments. Because first and second event durations were modulated independently in this experiment, we were able to model duration judgments as a function of both first and second event duration. The strongest version of our account — that event boundaries completely disrupt access to pre-boundary information, leading to reliance on only the second event — would predict that duration judgments would scale only with the duration of the second event, regardless of first event duration. We thus ran a linear model on the group-averaged data, predicting mean duration judgments as a linear combination of first and second event. Overall, this model proved an excellent fit to the data, R^2^= 0.99. There were significant contributions of both first (*β* = 0.59; *p* < 0.001) and second (*β* = 0.66; *p* < 0.001) event durations, though notably the weight on the second event was numerically higher. To assess the reliability of this model fit, as well as the difference in weights on the first versus second events, we performed bootstrap resampling across participants (Efron & Tibshirani, 1986). Briefly, we resampled our participants with replacement 1,000 times; with each iteration, we recomputed the group-averaged data as well as the model fits. We then derived an empirical *p* value by assessing for what proportion of resamples the weight on the second event was higher than that of the first. The weight on the second event was higher than the first in all 1,000 resamples (*p* < 0.001), indicating that although both first and second event durations influence duration judgments, participants overweight the duration of the second event. This finding is consistent with the interpretation that event boundaries lead to *reduced* (though not completely eliminated) accessibility of the first event, and thus an increased reliance on the post-boundary event.

### Experiment 3

Experiment 2 provided initial evidence that memory accessibility may contribute to duration judgments. Next, to approach this question in a complementary way, we examined whether abrupt changes (associated with flushing of working memory contents) were necessary to elicit boundary-related temporal compression. In a between-participants design, participants were tested on continuous and either “abrupt” boundary trials (with three color switches, equally spaced over the interval) or “gradual” boundary trials (in which the color smoothly morphed over the entire interval; **Figure 1G**). We hypothesized that if the context drifted gradually, rather than abruptly, over the course of the trial, we would observe a reduced effect of color change on duration judgments. Such gradual shifts may be associated with some forgetting of early information, but we would not necessarily expect a full “flushing” based on prior work showing that prior memories persist when change is gradual (Gershman et al., 2014).

## Methods

### Participants

Eighty individuals (62 female; age range 18-27; mean = 19.9) were recruited from New York University and the larger community and participated for either course credit or payment ($10/hr). Twenty individuals each were assigned to each of the four experimental groups (described below). Informed consent was obtained in a manner approved by the University Committee on Activities Involving Human Subjects at New York University. Post-hoc power analyses of Experiment 1 revealed an achieved power of 0.998, lending confidence to the choice of 20 per group.

### Stimuli

Stimuli were colored squares presented centrally on a mid-gray background. Each square was presented on-screen for a 0.5 – 5 seconds interval (sampled equally in increments of 0.5 seconds).

For all participants, for half of the trials, the color remained the same for the entire duration (continuous condition), as in Experiment 1. The other half of the trials were “boundary condition,” which differed by experimental group.

In the abrupt change groups, there were two color switches, which occurred one-third and two-thirds of the way through the total duration. To test whether the boundary duration compression effect was driven by the number of changes, or change of the final color specifically, we had an “ABC” group, in which the color switched to a new color at each switch, and an “ABA” group, in which the final segment of the trial returned to the color of the first segment.

In the gradual change groups, the color slowly morphed from the initial color to the final color without an abrupt boundary. Because both rate of color change and ending color (if rate of change was held constant) could be cues to duration, we included two groups: one which had a constant rate of change and different end colors for different durations (rate-constant), and one which had constant end colors and different rates of change for different durations (rate-changing). See **Supplemental Figure 2** for a depiction of the four conditions.

For the abrupt change groups, square stimuli were blue (R = 0, G = 0, B = 255), green (R = 0, G = 255, B = 0), and yellow (R = 260, G = 200, B = 80). For each participant, color triplets were randomly assigned such that the order of the three colors in the boundary condition was fixed. Each participant viewed each color/color-triplet twice for each time point.

For the gradual change groups, square stimuli morphed between red (R = 250, G = 0, B = 0) and blue; blue (R = 0, G = 0, B = 250) and red; and blue and green. In the rate changing group, the step size between colors per screen refresh was calculated separately for each timepoint such that the end color was always the same, e.g., starting at [250 0 0] and ending at [0 0 250]. In the rate constant group, the step size between colors per screen refresh was fixed, such that 1 RGB value was subtracted from the start color and one added to the end color for each refresh.

Pseudorandomization of trial sequences was conducted using the same procedure as in experiment 1. In total, the task consisted of 6 trials per condition and time point (120 trials total), broken up into two runs.

### Duration Judgment Task

The duration judgment task was identical to that described in Experiment 1.

#### Results

Again, we first assessed accuracy of participants’ duration judgments, and found that all participants exhibited a reliable correlation, mean z = 1.06, 95% CI = [0.99, 1.13], *t*(79) = 29.31, *p* < 0.001, *d* = 3.28. (**Supplemental Figure 1C)**.

Next, we assessed our key hypothesis that gradual, as opposed to abrupt, shifts in context will be associated with a reduced influence of event boundaries on temporal duration judgments. First, we examined each of the four experimental groups separately. Because there were no differences in the boundary effect for the two abrupt change and two gradual change groups, respectively (See Experiment 3 Stimuli Methods; **Supplemental Figure 2**), these groups were collapsed. In the abrupt change group, we robustly replicated the boundary compression effect (M = −0.17, 95% CI = [-0.24, −0.11]; *t*(39) = −5.14, *p* < 0.001, *d* = −0.81). In the gradual change group, we also replicated the effect (M = −0.08, 95% CI = [-0.15, −0.01]; *t*(39) = −2.30, *p* = 0.027, *d* = −0.36), though the effect was numerically weaker. Consistent with our hypothesis, an ANOVA revealed a marginal interaction between condition (continuous vs. boundary) and group (gradual vs. abrupt change), such that there was reduction in the boundary effect for the gradual group (F(1, 78) = 3.63, p = 0.060, *ηp^2^ =* 0.04; **Figure 1H**). These data suggest that abrupt changes in context (associated with decreased memory accessibility) result in stronger boundary effects, and provide further support that the extent of duration distortion across events is modulated by accessibility to the event representations.

### Experiment 4

Across the three behavioral experiments presented thus far, we extend prior work demonstrating that event boundaries reduce estimates of duration, and provide novel insight into *why* event boundaries may lead to reduced duration judgments; specifically, we provide evidence that such reductions may be due to decreased mnemonic accessibility. Next, we ran an fMRI experiment designed to assess how hippocampal pattern change within a single trial relates to temporal duration judgments.

#### Participants

Eighteen right-handed native English speakers (7 female; age range 22-34; mean = 27.1) were recruited from New York University and the larger community and participated for payment ($25/hr). We planned to recruit twenty individuals, as in Experiment 1 (in which achieved power was 0.998), but due to attrition only had eighteen usable participants. However, this experiment has more trials than the previous, and the neural analyses use a within-participant approach to brain-behavior correlations. Informed consent was obtained in a manner approved by the University Committee on Activities Involving Human Subjects at New York University. Due to equipment failure, the first eight out of fifteen runs were lost for one participant; thus, only the usable runs of this data set were included in analyses. Due to time constraints, the verbal temporal perception task and localizer task were not collected for two participants.

#### Stimuli

Stimuli were colored blue (R = 0, G = 0, B = 160), red (R = 160, G = 0, B = 0), and yellow (R = 195, G = 175, B = 35) squares presented centrally on a mid-gray background. In contrast to Experiments 1-3, each square was presented on-screen for a 2, 4, 6, or 8 second interval. For half of the trials, the color remained the same for the entire duration (continuous condition), and for the other half of the trials the color switched half-way through the total duration (boundary condition).

Each participant viewed each color or color-pair an equal number of times for each time point (here, ten times over the entire experiment). Color pairings and pseudorandomization of trial sequences was conducted using the same procedure as in Experiment 1.

During each delay period (see below), a colored noise mask was presented over the entire display. A single mask image was created by randomly sampling R, G, and B values independently for each pixel on a 13” MacBook Pro.

In total, the task consisted of 30 trials per condition and time point (240 trials total). These were broken into 15 fMRI runs of 16 trials each, such that each condition/duration combination was presented twice per run.

#### Duration Judgment Task

The duration judgment task was identical to that of Experiment 1, with the following changes. The timeline was bounded by 0 sec and 10 sec. Participants responded using an MRI-compatible trackball. Additionally, there was a variable delay interval (2, 4, or 6 seconds) between square presentation and time judgment, which was included to orthogonalize the fMRI regressors associated with the presentation and judgment periods. During this delay, a colored noise mask was presented over the entire screen to prevent any after-images that may be caused by a lingering percept of the square.

There was a variable inter-trial interval (4, 6, or 8 seconds) during which participants performed an odd/even task after the time judgment period. Each number was presented for a maximum of 1.9 seconds with an inter-stimulus interval of 0.1 seconds. Participants responded using the left and right buttons on the sides of trackball. The inter-trial periods were jittered in order to orthogonalize the fMRI regressors associated with the presentation and judgment periods, and this interval served as baseline in fMRI analyses.

#### Verbal duration judgment task

We additionally included a single behavioral run following completion of the fMRI study, in which participants – instead of indicating their responses on the number line – verbally reported how long they thought the square was on the screen for, in seconds (**Supplemental Figure 3C**,**D**). Participants were given no numerical bounds or constraints on how precise their responses had to be. No functional data were collected during this session. The goal of this additional run was to assess participants’ duration estimates in an unconstrained way (i.e., without giving them a bounding number line) and to verify that our effects could not be explained by idiosyncrasies in how individual participants use the number line.

#### Localizer

At the end of the fMRI session, participants engaged in a blocked color localizer in which each of the three colors was flashed repeatedly (0.9 seconds on, 0.1 seconds blank inter-stimulus interval) for 32 seconds. Each color was presented twice per run, and there were two runs of this task. The data from this run are not reported in this paper.

#### fMRI Parameters

Functional images were acquired on a Siemens Allegra head-only 3T scanner. Data were collected using an EPI pulse sequence (34 contiguous slices oriented parallel to the AC-PC axis; TR = 2000 ms; TE = 15 ms; flip angle = 82°; voxel size 3×3×3 mm). Additionally, a high-resolution T1-weighted anatomical scan (magnetization-prepared-rapid-acquisition gradient echo sequence, 1×1×1 mm) was obtained for each participant. During functional scans, stimuli were viewed through a mirror attached to the head coil.

#### Preprocessing of fMRI Data

fMRI data processing was carried out using FEAT (fMRI Expert Analysis Tool) Version 6.00, part of FSL (FMRIB’s Software Library, www.fmrib.ox.ac.uk/fsl). BOLD images were first skull-stripped using the Brain Extraction Tool (Smith, 2002). The first four volumes of each run were discarded to allow for T1 stabilization. Additionally, images underwent motion correction using MCFLIRT (Jenkinson et al., 2002), grand-mean intensity normalization by a single multiplicative factor, and high-pass temporal filtering (Gaussian-weighted least-squares straight line fitting, with sigma = 50.0s). No spatial smoothing was applied to the data. Each run was realigned to the final fMRI run of the session. Motion outliers were detected using FSL Motion Outliers; TRs containing motion outliers were excluded from pattern similarity analyses (see “Pattern Similarity Analysis” section below).

#### ROI Definition

The primary ROIs in this study were the hippocampus and visual cortex. However, given that many other regions have been implicated in temporal duration judgments (Buhusi & Meck, 2005; Lositsky et al., 2016), we also examined prefrontal regions, striatal regions, and the entorhinal cortex.

Anatomical hippocampal ROIs (left and right) were defined for each participant via FSL’s FIRST automated segmentation tool (Patenaude et al., 2011). We examined both the bilateral hippocampus, as well as each hemisphere separately. However, given prior findings that patterns specifically in the left hippocampus relate to memory and time estimates across boundaries (DuBrow & Davachi, 2014; Ezzyat & Davachi, 2014; Hsieh et al., 2014; Nielson et al., 2015), we hypothesized that our effects may be driven by the left hippocampus.

For our visual cortex ROI, we created a mask of V1 and V4. Specifically, we used probabilistic masks for V1 and hV4 (Wang et al., 2015); these masks were thresholded to 75%, concatenated together, and then transformed into each participant’s native space. This choice was motivated by prior work demonstrating patterns of activity in V1 and V4 carry information about color (Brouwer & Heeger, 2009).

Anatomical caudate and putamen ROIs were defined for each participant via FSL’s FIRST automated segmentation tool (Patenaude et al., 2011).

Prefrontal cortical regions were defined using the Harvard-Oxford Cortical Structural Atlas. Specifically, ROIs were created in standard space for Middle Frontal Gyrus; Inferior Frontal Gyrus, pars triangularis; and Inferior Frontal Gyrus, pars opercularis. Each of these three ROIs were then transformed into participants’ functional spaces and thresholded at 75%. The Inferior Frontal Gyrus, pars triangularis and Inferior Frontal Gyrus, pars opercularis were then concatenated together to create one IFG ROI.

Entorhinal cortex was manually segmented on every participant’s T1-weighted anatomical scans, using published anatomical landmarks (Insausti et al., 1998; Pruessner et al., 2002).

#### Pattern Similarity Analysis

Pattern similarity analyses were performed on 8 second trials only to maximize separation between the hemodynamic responses to the beginning and end of each trial. The data were modeled via an approach adapted from Turner et al (2012). This approach is well suited for within-trial analyses, as it uses a multi-parameter single-trial GLM, which allows for the un-mixing of temporally adjacent BOLD responses. Because our stimuli were temporally extended, we extracted the activation pattern across voxels for TRs 2-7 (4-14 seconds) after the true stimulus onset. A design matrix was constructed in which each of these “trial-related” TRs was a separate regressor of interest; in addition, separate nuisance regressors of no-interest included separate TRs for every other trial collapsed across each condition/color combination, as well as TRs from the ITI and delay periods. This design matrix was regressed against the activation of each voxel in each ROI to compute a separate beta estimate for each TR of the trial. To avoid making strong assumptions about where the true onset, midpoint, and offset of each trial fall, we estimated the beginning and end of the stimulus period by averaging the first two and last two betas, respectively. Specifically, from trial onset, the second and third TRs (corresponding to approximately 0-2 seconds after stimulus onset) were averaged as an estimate of the “beginning” and the fifth and sixth TRs (corresponding to approximately 6-8 seconds after stimulus onset) were averaged as an estimate of the “end” (see Fig 2).

**Figure 2.**
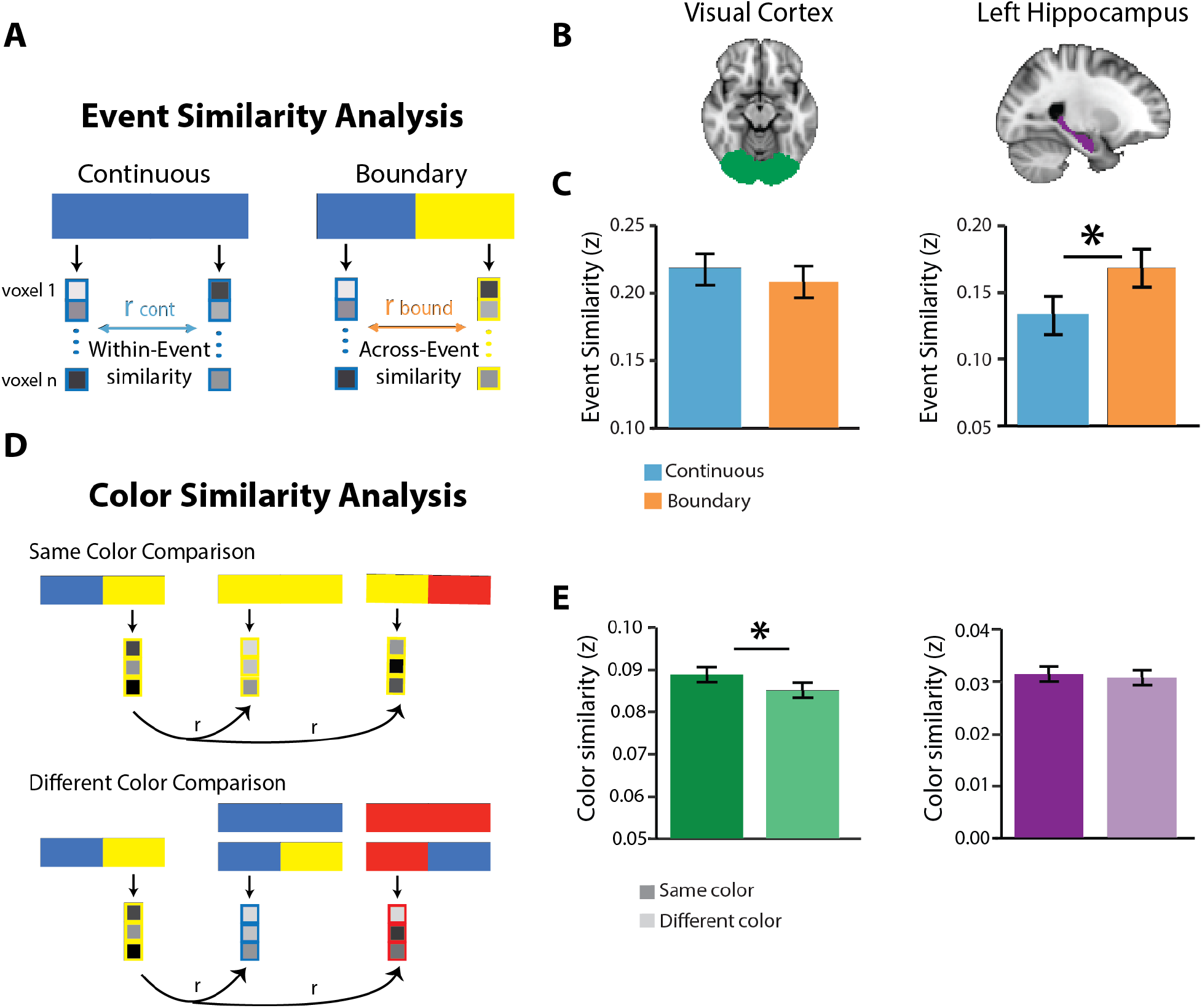
Pattern Similarity Approaches. A) Event similarity was computed as the fisher-transformed Pearson correlation between the estimated evoked activity at the beginning of an 8 second trial and the estimated evoked activity at the end of that 8 second trial. Correlations were computed separately for continuous sand boundary trials. B) Visual cortex did not show differential event similarity by condition. C) The left hippocampus exhibited greater event similarity across boundary trials than across continuous trials. D) Color similarity was computed as the fisher-transformed Pearson correlation between the estimated evoked activity at the end of an 8 second trial, to the estimated evoked activity at the beginning of another trial that shares the same color. As a comparison, “different color” trials were computed as the correlation between the patterns at the end of one trial to the beginning of all other trials which did not share its color. E) The visual cortex exhibited greater similarity for trials of the same color than trials of a different color. F) Left hippocampus did not show differential similarity as a function of trial color. * p < .05, two-tailed. Error bars denote the within-participant standard error of the mean.

TRs containing a motion outlier were excluded from subsequent analyses; furthermore, if there was more than one motion outlier within the “trial-related” TRs, the entire trial was excluded from subsequent analyses. On average, 5 (SD = 3.33) individual timepoints per participant were excluded from our analysis. Further, an average of 2.83 (SD = 1.34) trials per participant from our analysis were excluded due to there being more than one motion outlier in a given trial.

#### Event Similarity

Event similarity was computed by correlating the patterns corresponding to the “beginning” and “end” of every trial separately. Pearson correlation values were Fisher-transformed and averaged for the two conditions to produce a “similarity” score.

#### Event Similarity Predicting Behavior

To assess whether within-trial event similarity in visual cortex and hippocampus influence duration judgments, we performed a mixed-effects analysis implemented via the lme4 package in R (http://cran.r-project.org/web/packages/lme4). To account for individual differences in number line usage, we z-scored participants’ duration judgments across all trials. Mixed-effects linear models were run, and significance was assessed using iterative model comparisons, which resulted in chi-squared values and corresponding *p*-values. Normalized duration judgment was the dependent measure; visual cortical similarity, hippocampal similarity, and condition, as well as interactions between them, were included as fixed-effect predictors. Our general approach consisted of running the simplest form of the model, and then only including significant predictors from these simple models in subsequent models. Model comparisons were also used to determine which variables were included as by-participant random factors; as a result of these comparisons, the condition factor was included as a by-participant random effect, such that a slope and intercept were calculated for each participant.

#### Color Similarity

To isolate responses to color, irrespective of temporal position information, we assessed similarity of the “end” of a boundary trial to the “beginning” of all other trials, conditionalized on whether the color at the end of the trial of interest was the same or different than the color at the beginning of the other trials. This across-trial color similarity metric was computed as the Fisher transformation of the Pearson correlation between the “end” of the trial of interest and the “beginning” of every other trial separately. Trials from the same fMRI run were excluded due to temporal autocorrelation. For each trial, the mean of the similarity values to all other trials was computed. The mean color similarity for “same” and “different” color comparisons was then calculated.

## Results

### Behavior

First, we again assessed accuracy of participants’ duration judgments, and found that all participants exhibited a reliable correlation, mean z = 1.08, 95% CI = [0.96, 1.20], *t*(17) = 18.98, *p* < 0.001, *d* = 4.47 (**Supplemental Figure 1D)**.

Next, we sought to replicate the finding from Experiment 1, with longer (8 sec) trials in the fMRI study (**Figure 1I**). Indeed, we found that boundary trials were judged as significantly shorter than continuous trials (M = −0.31, 95% CI = [-0.61, −0.02]), *t*(17) = −2.26, *p* = 0.037, *d* = −0.53 (**Figure 1J**). We additionally ran this analysis for all trials, both within duration and collapsed across, and found a similar pattern of results **(Supplemental Figure 3A**,**B)**. We also separately examined data from the additional run of the task, in which participants responded verbally rather than with the number line. We found a comparable boundary effect even within that single run **(Supplemental Figure 3C)**, and an across-participant correlation between the boundary effect across the whole fMRI task (15 runs) and the single verbal run **(Supplemental Figure 3D)**. This additional run demonstrates the robustness of the results, providing evidence that these effects can be observed within a single run, and with a different, unconstrained response medium.

Given that in this experiment, unlike Experiments 1-3, there was a brief delay between the square presentation and the duration judgment, we also verified that the delay did not have an influence on duration judgments. Indeed, across all trials, although there was a marginal main effect of condition (*F*(1, 17) = 3.86, *p* = 0.066, *ηp^2^ =* 0.19), there was no main effect of ISI on duration judgments (*F*(2, 34) = 0.012, *p* = 0.988, *ηp^2^ =* 0.00), nor an interaction between condition and ISI duration (*F*(2, 34) = 0.52, *p* = 0.600, *ηp^2^ =* 0.03). The same pattern was true for the 8 sec trials only: although there was a marginal main effect of condition (*F*(1, 17) = 4.18, *p* = 0.057, *ηp^2^ =* 0.20), there was no main effect of ISI on duration judgments (*F*(2, 34) = 0.42, *p* = 0.661, *ηp^2^ =* 0.02), nor an interaction between condition and ISI duration (*F*(2, 34) = 0.21, *p* = 0.812, *ηp^2^ =* 0.01).

### Event Similarity in the Hippocampus

In Experiment 4, our primary interest was how hippocampal representational change within individual trials is related to behavioral duration judgments. To first examine the influence of event structure on neural similarity measures, we examined differences in similarity across individual 8-second trials. Specifically, we extracted the pattern of activity across voxels at the beginning of the trial and at the end of the trial, and correlated those two patterns to measure the representational change across the interval. This was done separately for continuous and boundary trials (**Figure 2A**). Thus, in the continuous condition, we examined changes in patterns of activity from the beginning to the end of a single event, whereas in the boundary condition, we probed changes in patterns across two events. Importantly, however, the interval length in the two conditions was matched. This analysis allows us to examine representations that occur at the event-level: if a region represents events as a period of neural stability (DuBrow and Davachi, 2014; Baldassano et al., 2017; Ezzyat & Davachi, 2014, 2021), then we would expect to see greater similarity for the continuous condition, which contains one event, versus the boundary condition, which contains two. However, if a region is sensitive to the temporal position information within an interval (e.g., the start of an event), then we may expect to see greater similarity in the boundary condition, akin to a “resetting” of the neural population at a boundary (Levy, 1989; Wallenstein et al., 1998; Ben-Yakov et al., 2013; Terada et al., 2017; Ben-Yakov & Henson, 2018; Sun et al., 2020).

We focused on two regions of interest: our primary region of interest, the hippocampus, and the visual cortex (V1/V4) as a control region (See ROI Definition Methods; **Figure 2B**). Specifically, we hypothesized that pattern similarity in the visual cortex would decrease in the boundary condition at a color switch, since it should be sensitive to perceptual similarity. Contrary to our hypothesis, we did not find differences in similarity by condition in the visual cortex, (M = 0.01, 95% CI = [-0.02, 0.03], *t*(17) = 0.79, *p* = 0.439, *d =* 0.19 **(Figure 2C, left)**. In the bilateral hippocampus, however, there was marginally greater event similarity for the boundary condition, relative to continuous, M = −0.03, 95% CI = [-0.06, 0.00], *t*(17) = −1.88, *p* = 0.077, *d =* −0.44 (**Supplemental Figure 4A, left**). This effect was driven by the left hippocampus, which exhibited significantly greater event similarity for the boundary condition, M = −0.04, 95% CI = [-0.07, −0.01], *t*(17) = −2.50, *p* = 0.023, *d* = −0.59. (**Figure 2C, right**). In contrast, the right hippocampus exhibited no difference in event similarity as a function of condition, M = −0.02, 95% CI = [-0.06, 0.02], *t*(17) = −1.21, *p* = 0.244, *d* = −0.28 (**Supplemental Figure 4A, right**).

To assess whether the effects observed in the left hippocampus significantly differed from the pattern observed in visual cortex, we further conducted an ANOVA examining pattern similarity as a function of both ROI and condition. There was no main effect of condition on pattern similarity, *F*(1, 17) = 1.81, *p* = 0.196, *ηp^2^ =* 0.10, though there was a main effect of ROI on pattern similarity, *F*(1, 17) = 11.97, *p* = 0.003, *ηp^2^ =* 0.41. Critically, there was a significant interaction between condition and ROI, *F*(1, 17) = 6.88, *p* = 0.018, *ηp^2^ =* 0.29, indicating that visual cortex and hippocampus exhibited distinct effects.

### Color Similarity in Visual Cortex

The event similarity analysis provided evidence that left hippocampal patterns, but not visual cortical patterns, were modulated by the presence of an event boundary. To ensure that the dissociation between hippocampus and visual cortex was not just due to differences in task sensitivity, we next asked whether patterns of brain activity were modulated by color, irrespective of events. To isolate color representations irrespective of event representations, we correlated the activity patterns from the second event of one (boundary) trial to the first event of all other trials (**Figure 2D**). We then sorted the data by whether the first and second events were the same color or different colors, and examined differences in pattern similarity as a function of color match. In the visual cortex, we found that pattern similarity was greater for two events of the same color, compared to two events of different colors (M = 0.004, 95% CI = [0.00, 0.001]), *t*(17) = 2.14, *p* = 0.047, *d* = 0.51 (**Figure 2E, left**), indicating visual cortex was sensitive to color information in this task. In contrast, the left hippocampus, which exhibited sensitivity to event structure, showed no such sensitivity to color, (M = 0.00, 95% CI = [-0.00, 0.00]), *t*(17) = 0.48, *p* = 0.637, *d* = 0.11 (**Figure 2E, right**). The right hippocampus (M = −0.001; 95% CI = [-0.004, 0.002], *t*(17) = −0.81, *p =* 0.430, *d* = −0.19) and bilateral hippocampus (M = 0.00; 95% CI = [-0.003, 0.002], *t*(17) = −0.37, *p* = .719, *d* = −0.09) exhibited similar null results.

To again assess whether the effects observed in the visual cortex significantly differed from the hippocampus, we further conducted an ANOVA examining pattern similarity as a function of both ROI and condition (same vs different color). We ran this analysis in the left hippocampus, as this was the ROI which exhibited sensitivity to event structure, but the results are comparable in the right and bilateral hippocampus. There was no main effect of condition on pattern similarity, *F*(1, 17) = 2.73, *p* = 0.117, *ηp^2^ =* 0.14, though there was a main effect of ROI on pattern similarity, *F*(1, 17) = 40.52, *p* < 0.001, *ηp^2^ =* 0.70. There was a marginal interaction between condition and ROI, *F*(1, 17) = 3.28, *p* = 0.088, *ηp^2^ =* 0.16.

### Relationship between event dissimilarity and duration judgments

Finally, we asked whether pattern change over the course of a single 8-second trial could explain variance in duration judgments across trials. To examine how neural pattern similarity influences duration judgments, we ran a series of mixed-effects linear regressions. Specifically, we tested whether event dissimilarity (1 - ‘Event Similarity’ described above) in hippocampus and in visual cortex predict duration judgments above and beyond the influence of event boundaries, as well as whether they interacted with condition.

First, we tested a model predicting duration judgment from both condition and hippocampal pattern dissimilarity, and we found a main effect of bilateral hippocampal dissimilarity (*β* = 0.29; *χ*^*2*^(1)= 4.67, *p* = 0.031; **Supplemental Figure 4B, left**), such that greater dissimilarity — or more pattern change — in the hippocampus predicted longer duration judgments. When examining this effect separately for each hemisphere, we found that right hippocampal dissimilarity did not predict duration judgment (*β* = 0.14; *χ*^*2*^(1) = 1.17, *p* = 0.280; **Supplemental Figure 4B, right**), whereas (consistent with the event similarity result) left hippocampal dissimilarity did (*β* = 0.40; *χ*^*2*^(1) = 9.68, *p* = 0.002; **Figure 3A**). When including both the right and left hippocampal pattern dissimilarity (in addition to condition) in a model predicting duration judgment, the left hippocampus reliably predicted duration judgment (*β* = 0.57; *χ*^*2*^(1)= 10.46, *p* = 0.001), whereas the right hippocampus did not (*β* = −0.25; *χ*^*2*^(1)= 2.04, *p* = 0.153). Thus, in all subsequent models we will only consider the left hippocampus.

There was no significant interaction between left hippocampal dissimilarity and condition (*χ*^*2*^(1) = 2.48, *p* = 0.115). Although this effect of hippocampal dissimilarity on duration judgments was only significant in the continuous condition (*β* = 0.59; *χ*^*2*^(1)= 9.10, *p* = 0.003), it was numerically in the same direction in the boundary condition (*β* = 0.21; *χ*^*2*^(1)= 1.59, *p* = 0.207).

Next, we constructed a model predicting duration judgment from visual cortical dissimilarity, in addition to condition. The model revealed no main effect of visual cortex dissimilarity on duration judgments (*β* = 0.17; *χ*^*2*^(1)= 1.33, *p* = 0.249; **Figure 3B**). However, a model predicting duration judgment from the main effect of condition as well as an interaction between visual cortical dissimilarity and condition, revealed a significant interaction between visual cortex dissimilarity and condition (*χ*^*2*^(2) = 6.37, *p* = 0.041). Specifically, visual cortical dissimilarity predicted duration judgments in the continuous condition (*β* = 0.48; *χ*^*2*^(1)= 5.08, *p* = 0.024) but not the boundary condition (*β* = −0.17; *χ*^*2*^(1)= 0.78, *p* = 0.378).

**Figure 3.**
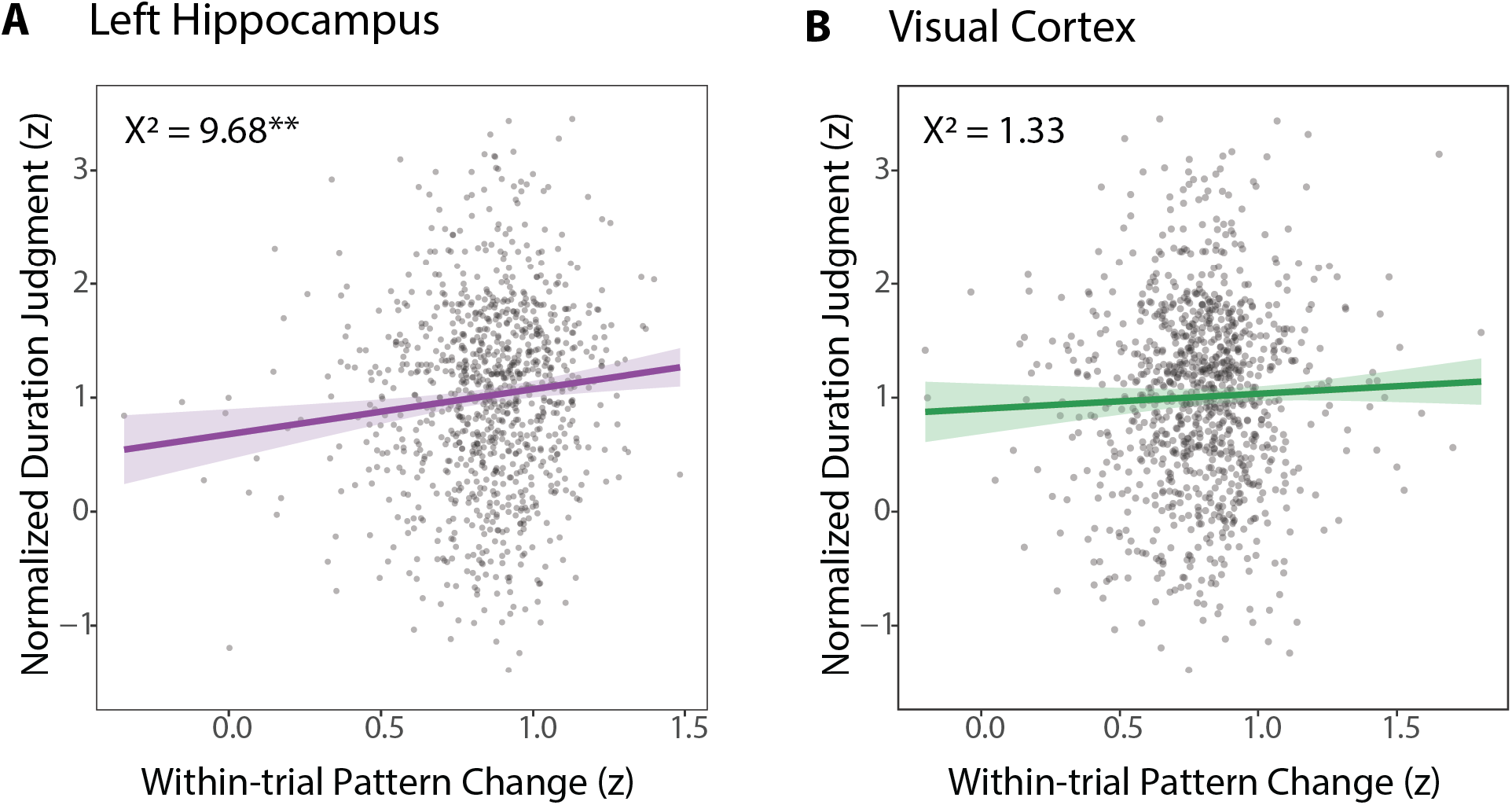
Results of mixed-effects linear model predicting time judgment by event dissimilarity. A) Event dissimilarity in the left hippocampus predicted duration judgments. Plotted from a model predicting time judgment as a function of: condition, a main effect of a main effect of hippocampal dissimilarity, and a by-participant random effect of condition. B) Event dissimilarity in visual cortex did not predict time judgment across conditions. Plotted from a model predicting time judgment as a function of: condition, a main effect of a main effect of visual cortical dissimilarity, and a by-participant random effect of condition.

Given that both the left hippocampus and visual cortex dissimilarity exhibited relations with duration judgments in the continuous condition, we asked whether they contribute independent variance. In a model predicting duration judgment in the continuous condition as a function of left hippocampal and visual cortical dissimilarity, we found that the hippocampus significantly contributed to duration judgment (*β* = 0.49; *χ*^*2*^(1)= 4.91, *p* = 0.027), but the visual cortex did not (*β* = 0.23; *χ*^*2*^(1)= 0.89, *p* = 0.345).

Given that many other brain regions have been implicated in time perception at this timescale (Buhusi & Meck, 2005; Lositsky et al., 2016), we exploratorily performed this analysis in other regions. Specifically, we assessed whether pattern dissimilarity in the caudate, putamen, middle frontal gyrus (MFG), inferior frontal gyrus (IFG), and entorhinal cortex predicted duration judgments. We did not find main effects of dissimilarity on duration judgments, nor interactions with conditions, in any of these regions (caudate: main effect, *β* = −0.12, *χ*^*2*^(1) =0.63, *p* = 0.427, interaction, *χ*^*2*^(2) =0.89, *p* = 0.639; putamen: main effect, *β* = 0.24, *χ*^*2*^(1) =0.68, *p* = 0.411, interaction, *χ*^*2*^(2) =0.70, *p* = 0.703; MFG: *β* = 0.14, main effect, *χ*^*2*^(1) =0.85, *p* = 0.355, interaction, *χ*^*2*^(2) =1.42, *p* = 0.493; IFG: main effect, *β* = 0.27, *χ*^*2*^(1) =3.38, *p* = 0.066, interaction, *χ*^*2*^(2) = 3.83, *p* = 0.147; entorhinal cortex: main effect, *β* = 0.052, *χ*^*2*^(1) =0.43, *p* = 0.510, interaction, *χ*^*2*^(2) = 0.50, *p* = 0.778).

## Discussion

Using event segmentation to manipulate mnemonic content, we found that duration judgments are influenced by the accessibility of mnemonic representations. After demonstrating that event boundaries lead to reduced duration judgments (Experiment 1), we found that duration judgments scale (to a limit) with the duration of the *most recent* event (Experiment 2) and that this reduction is attenuated when there is a gradual change in context that may maintain access to previous mnemonic information (Experiment 3). Lastly, trial-by-trial neural pattern change in the left hippocampus predicted longer duration judgments (Experiment 4).

### Relation to studies of event boundaries in short- and long-term duration judgments

Under a context change account of duration judgments, in which change is used to infer the passage of time, one might expect boundary trials to be judged as longer than continuous trials (Poynter, 1983; Zakay et al., 1994) because the context across two events differs more than the context within a single event. Yet, our results show the opposite: intervals with a boundary are estimated as *shorter* than equivalent, continuous intervals. Rather, our results point to a memory-based account of duration judgments, in which active mnemonic information is used to infer the passage of time: event boundaries disrupt access to pre-boundary information (Zwaan, 1996), leading to decreased time judgments despite greater context change. Notably, we do not propose that the pre-boundary information is completely lost, but rather its access is attenuated; such an interpretation is supported by the results of Experiment 2, which finds some influence of the first event duration on duration judgments. However, although our experiments were designed to *manipulate* accessibility to pre-boundary information, none provide a direct, empirical measure of the accessibility the pre-boundary information. Future work employing paradigms which simultaneously measure both memory accessibility/content, as well as duration judgments, could provide a more direct link.

In contrast to the current results, work at longer time scales shows precisely the opposite: intervals spanning a boundary are remembered as *longer* than equivalent continuous intervals, and duration judgments scale with the number of events (Ezzyat & Davachi, 2014; Faber & Gennari, 2015; Lositsky et al., 2016; Faber & Gennari, 2017). This discrepancy can be explained by the memory-based account, as event boundaries may affect mnemonic content differently in working versus long-term memory. That is, if event boundaries disrupt memory at encoding, there is less information in working memory, leading to an in-the-moment compression. However, event boundaries are often better remembered (Polyn et al., 2009; Swallow et al., 2009; Heusser et al., 2018; Rouhani et al., 2020), and may serve as anchor points to within-event content in long-term memory (DuBrow & Davachi, 2016; Michelmann et al., 2019; Shin & DuBrow, 2021). If intervals containing event boundaries have greater mnemonic content in long-term memory, this may lead to the inference that more time has passed. That said, future work using more directly comparable paradigms across short- and long-term judgments will be critical to tease apart these interpretations.

Notably, the differential results across short- and long-term paradigms cannot be explained by a key distinction in the interval timing literature: retrospective versus prospective timing (Block et al., 2018). In prospective paradigms, participants know in advance that they will be asked to report temporal information, and thus can attend to time. In retrospective paradigms, in contrast, participants are unaware that they will be asked to report temporal duration information. In long-term memory, event boundaries exert a similar influence on temporal memory, regardless of whether the paradigm is prospective (Ezzyat & Davachi, 2014; Faber & Gennari, 2017) or retrospective (Lositsky et al., 2016; Faber & Gennari, 2015). The current study, as well as other short-term studies (e.g., Bangert et al., 2019; Yousif & Scholl, 2019), employed a prospective design. Although — given converging results across prospective and retrospective in long-term memory studies — we do not think that a retrospective design would reverse the behavioral result, future work employing single-trial versions of the experiment could directly test this.

### Event representations in the hippocampus

We found increased hippocampal pattern similarity across trials containing a boundary, relative to continuous trials. Given that events are a period of contextual stability (DuBrow et al., 2017), this finding may seem counterintuitive. However, our finding converges with theoretical and empirical work that some hippocampal neurons are sensitive to specific temporal positions within an event (Levy, 1989; Wallenstein et al., 1998; Ginther et al., 2011; Terada et al., 2017; Sun et al., 2020). If there are neurons that code for the “beginnings” of events, an event boundary might recruit the same neural population as the beginning of the trial, yielding greater similarity across the two events; in contrast, for a continuous trial, the hippocampal representations would continue to grow dissimilar over time.

Alternatively, increased pattern similarity across boundaries may reflect enhanced pattern separation within events: such separation could orthogonalize experiences within an event and reduce within-context interference (Benear et al., 2020; but see Hsieh et al., 2014; Milivojevic et al., 2016). To distinguish between a position sensitivity vs. pattern separation account, future work with higher temporal precision could examine whether hippocampal pattern similarity across boundaries is driven by a change occurring specifically at the moment of the boundary, or continuous pattern change across the interval.

We found no evidence for event sensitivity in visual cortex. This effect was surprising, given findings that visual areas show greater pattern similarity within, versus across, events (e.g., Baldassano et al., 2017; Ezzyat & Davachi, 2021). However, although we may expect visual cortex to exhibit enhanced similarity for visually similar information, there may be greater neural adaptation in the continuous condition. Because our stimuli were temporally extended colored squares and BOLD signal in visual cortex becomes reduced for prolonged stimuli (Krekelberg et al., 2006), it is possible that adaptation countered any sensitivity to shared visual information.

### The role of the hippocampus in tracking time

Our finding that left hippocampal pattern change correlates with subjective duration judgments is consistent with its proposed role in representing time (Eichenbaum, 2013). In humans, left hippocampal pattern stability has been related to remembered temporal proximity, such that events remembered as “farther” apart show greater pattern dissimilarity (Ezzyat & Davachi, 2014; Nielson et al., 2015), consistent with our current results. Together, these findings suggest that temporal dissimilarity in hippocampal representations may represent subjective duration, across different timescales of both working and long-term memory — despite the fact that event boundaries have opposing effects on behavior at these two timescales. However, future work directly comparing the hippocampal patterns predicting temporal memory across short- and long-timescales (e.g., in the same task or same participants) is needed to more comprehensively understand how the left hippocampus may dually support these behaviors.

Notably, one study found that sequences (items + time), but not time alone, could be decoded from hippocampus (Thavabalasingam et al., 2019). Although these findings suggest that the hippocampus may not be able to represent temporal information in isolation, it is notable that they attempted to decode objective duration. In the current study we find that hippocampal patterns are related to *subjective* estimates of time, raising the possibility that the hippocampus codes for subjective, but not objective, duration.

That said, although we find that hippocampal patterns correlate with duration judgments, hippocampal damage does not consistently affect subjective duration judgments (e.g., Melgire et al., 2005; Noulhiane et al., 2007), especially at short durations (Jacobs et al., 2013; Palombo et al., 2016). However, the exact time scale at which processing becomes hippocampal-dependent is the subject of ongoing research, may be shorter than previously established (Sabariego et al., 2020), and may depend on the context of the task (Palombo et al., 2019). Our findings contribute to growing evidence that the hippocampus contributes to temporal processing at short timescales.

Prior work has identified a similar role for entorhinal cortex in representing time (e.g., Lositsky et al., 2016; Umbach et al., 2020). Surprisingly, we did not find that entorhinal patterns predicted duration judgments, though we note that our fMRI sequences were not optimized to collect signal from this region, which is prone to signal drop-out.

### Conclusions

One theoretical position is that brains, in fact, do not sense time; just because activity in a brain region is correlated with the passage of time does not mean that the region is clocking time per se (Ezzyat & Davachi, 2014; Davachi & DuBrow, 2015; Buzsáki & Llinás, 2017). Although here we report neural measurements that correlate with subjective duration judgments, we suggest that our results are a measure of event *memory* used to infer the passage of time. Our findings thus raise questions about the extent to which our sense of time arises from pure timing signals in the brain, or to what extent time is a product of — or reconstruction from — our memories.

## Acknowledgments

At the initial submission, we dedicated this work to Warren Meck and Howard Eichenbaum, pioneers in the study of time and the brain but, more importantly, both incredible people. Over the course of revising this paper, we tragically lost another pioneer: our dear friend, student, mentor and co-author Sarah DuBrow. Sarah was incredibly kind, sharp, and fun, and we miss her immensely. We hope she would be proud of the final paper.

We thank Emily Cowan and Alexa Tompary for helpful conversations, and Sami Yousif for feedback on the manuscript.

## Open Practices Statement

None of the studies reported in this article are pre-registered. The data for the study are available at the following OSF link: https://osf.io/n5pkb/?view_only=a1d4f756484e4f649fb25a9a166f0cbf

