## Supplemental Figures 1-4 for "Mnemonic content and hippocampal patterns shape judgments of time"

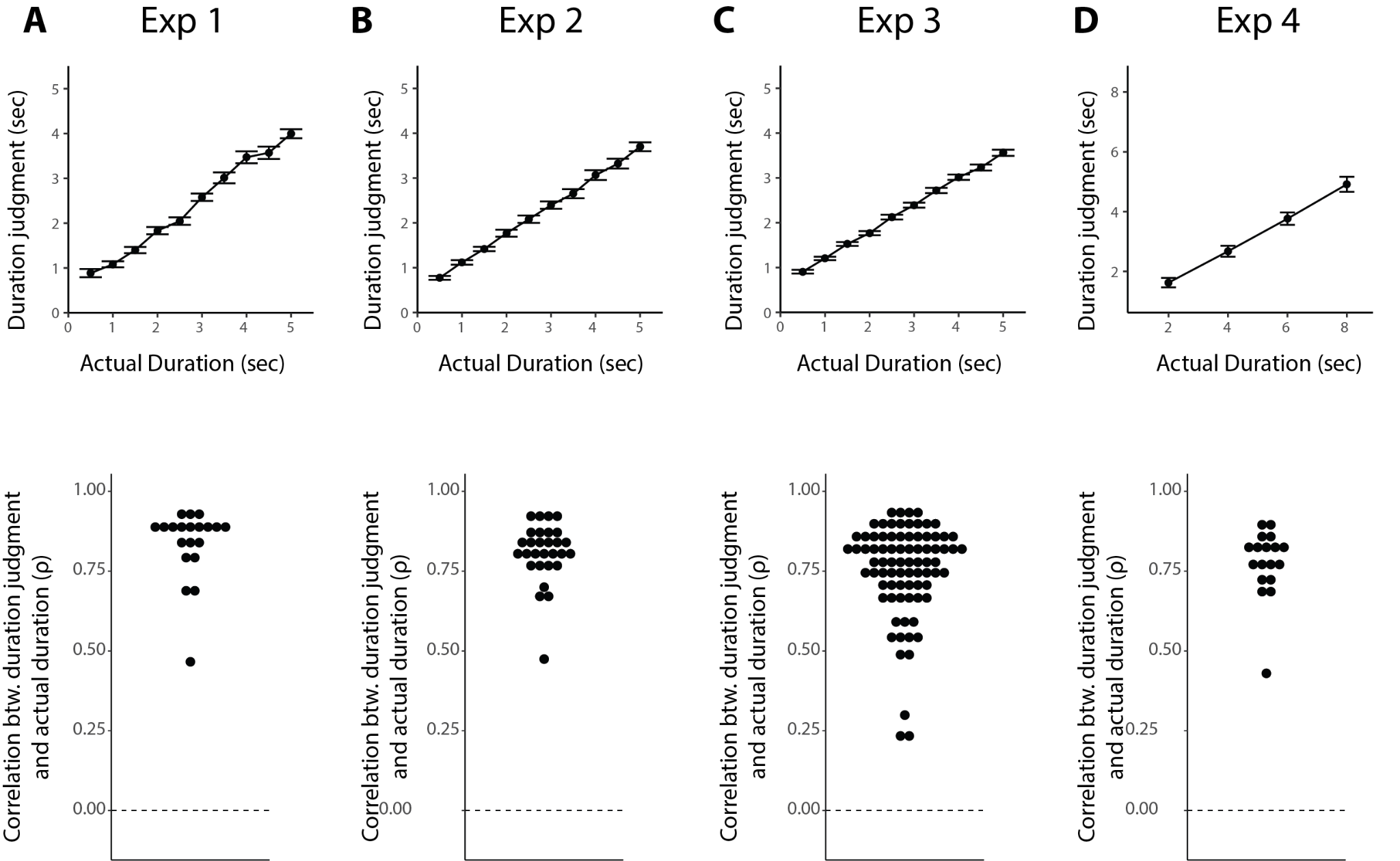

**Supplementary Figure 1. Correlations between duration judgment and actual duration.** A-D) Top: Mean duration judgments as a function of true durations for Experiments 1-4. Error bars represent standard error of the mean across participants. Bottom: within-participant Spearman Rank Correlations (rho) between the true duration and judged duration across all trials. Each dot is the correlation for one participant. All participants exhibited correlations greater than 0, and significantly above chance relative to a null distribution generated from their own responses.

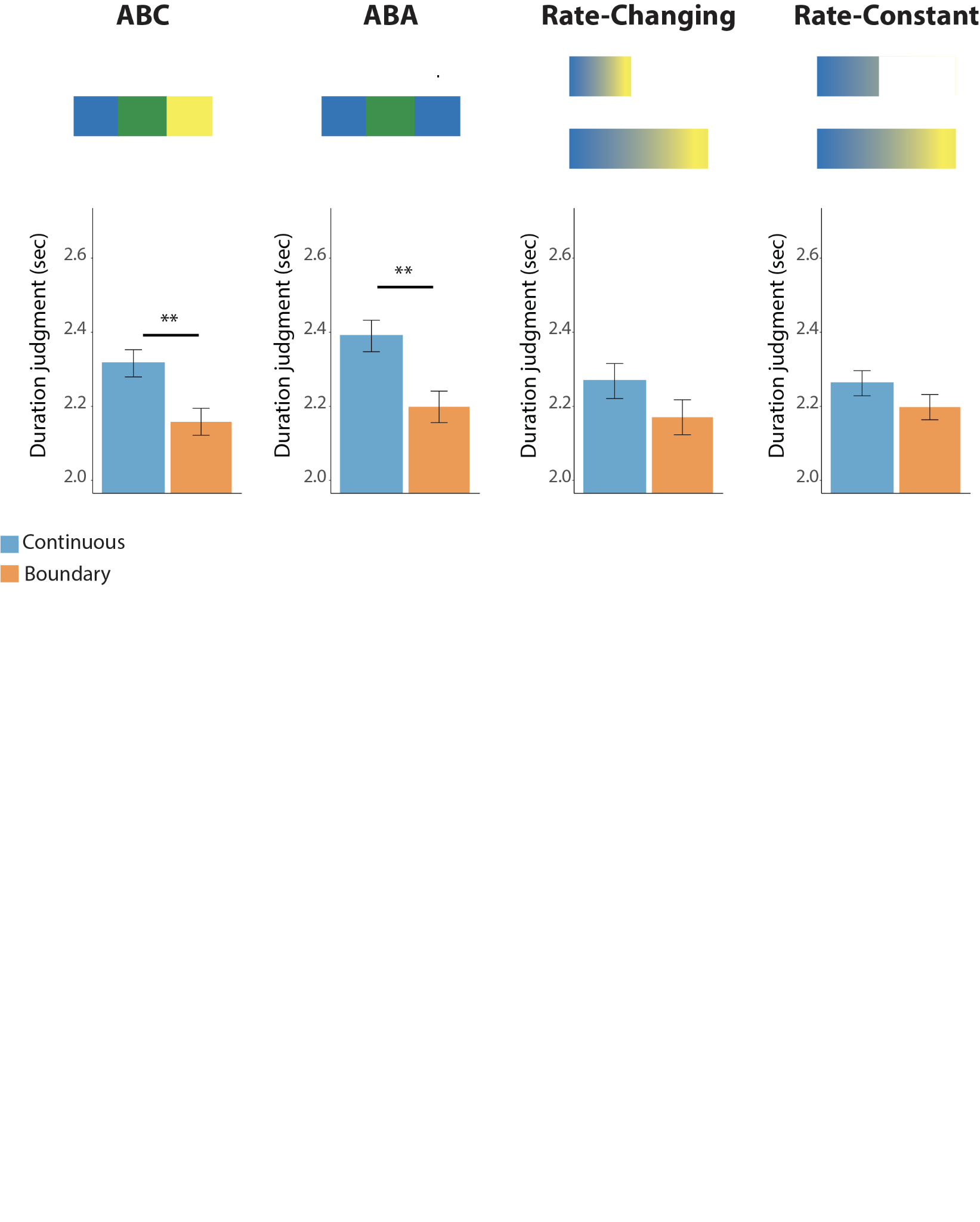

**Supplementary Figure 2. Depiction of the four conditions and results for Experiment 3.** ABC: three distinct colors were shown, each switching one-third of the way through the total duration. Boundary trials were judged to be significantly shorter than continuous trials, M = -0.16, 95% CI = [-0.25, -0.06], *t*(19) = 3.51 *p* = 0.002, *d* = -0.79. ABA: three colors were shown, with the third color being the same as the first. Each color switch occurred one-third of the way through the total duration. Boundary trials were judged to be significantly shorter than continuous trials, M = -0.19, 95% CI = [-0.30, -0.08], *t*(19) = 3.68, *p* = 0.002, *d* = -0.82. Rate-changing: the color drifted gradually over the entire duration. The rate of change differed across intervals such that the end color was held constant across the different durations. Boundary trials were judged to be non-significantly shorter than continuous trials, M = -0.10, 95% CI = [-0.22, 0.02], *t*(19) = 1.69, *p* = 0.11, *d* = -0.38. Rate-constant: the color drifted gradually over the entire duration. The rate of change was held constant across intervals such that the end color differed as a function of duration (more morphing for longer durations). Boundary trials were judged to be non-significantly shorter than continuous trials, M = -0.06, 95% CI = [-.15, 0.02], *t*(19) = 1.55, *p* = 0.14, *d* = -0.35. ** p < .005, two-tailed. Error bars denote the within-participant standard error of the mean.

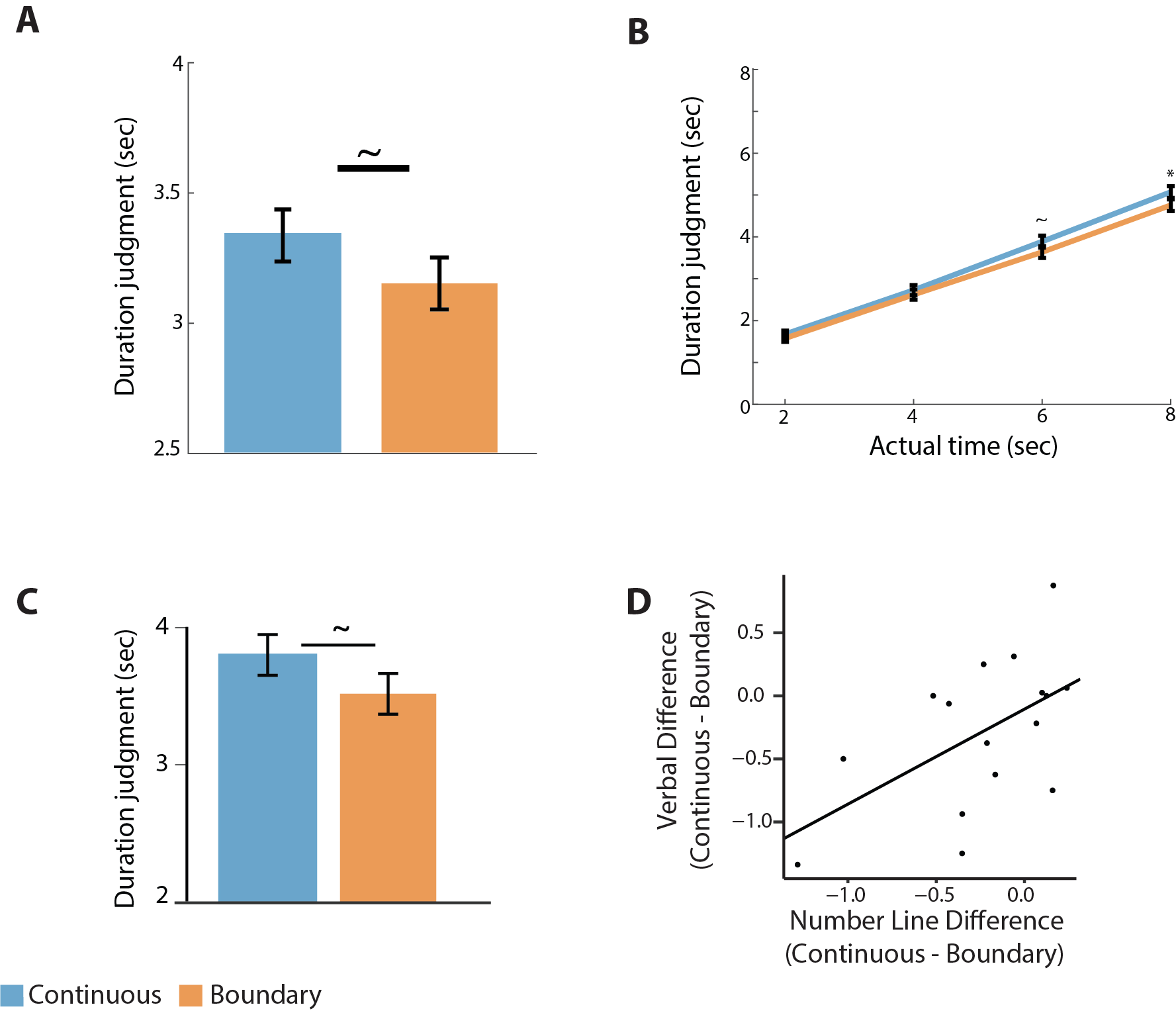

**Supplementary Figure 3. Additional behavioral results for Experiment 4.** A) Duration judgment as a function of condition, collapsed across all durations. Boundary trials were judged to be marginally shorter than continuous trials, M = -0.19, 95% CI = [-.40, 0.02], *t*(17) = 1.93, *p* = 0.070, *d* = -0.46. B) Duration judgment as a function of condition, separately for each duration. C) Duration judgments on the verbal duration judgment task, collapsed across duration. Boundary trials were judged to be marginally shorter than continuous trials, M = -0.28, 95% CI = [-0.60, 0.03], *t*(15) = 1.931 *p* = 0.075, *d* = -0.48. D) Across-participants correlation between the behavioral effects (duration judgment for continuous trials - duration judgment for boundary trials) on the verbal and number line duration judgment tasks. Each dot is a single participant. There was a reliable correlation in the boundary effects across the fMRI runs (number line difference) and subsequent verbal run (verbal difference); *r*(14) = 0.55, *p* = 0.028. ~p < 0.10; * p < 0.05. Error bars denote the within-participant standard error of the mean.

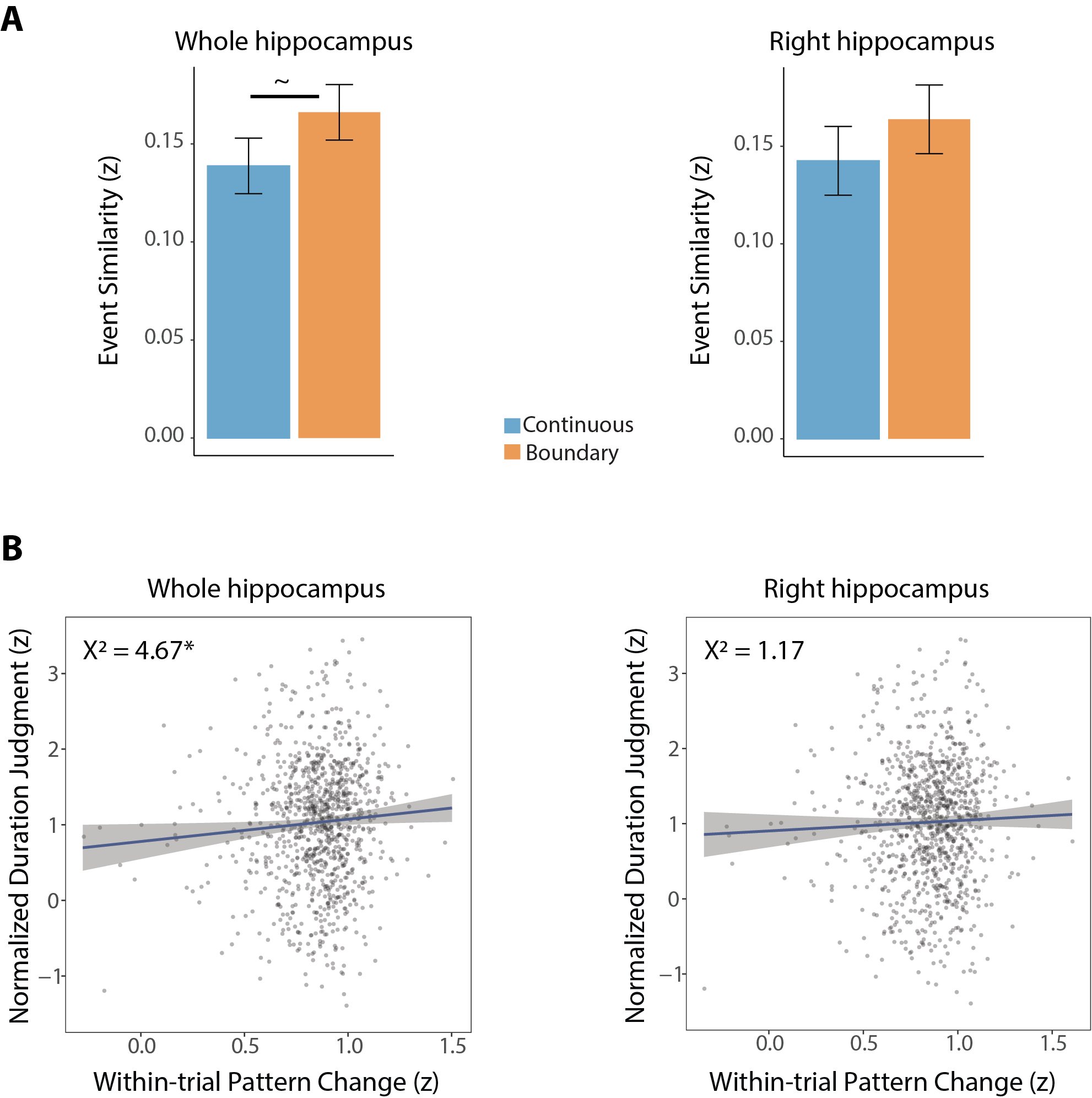

**Supplementary Figure 4. Whole and right hippocampus results.** A) Event similarity as a function of condition, in the bilateral hippocampus (left) and right hippocampus (right). In the bilateral hippocampus, there was marginally greater event similarity in the boundary condition, relative to the continuous condition, M = -0.03, 95% CI = [-0.06, 0.00], *t*(17) = -1.88, *p* = 0.077, *d =* -0.44. In the right hippocampus, there was no difference in event similarity as a function of condition, M = -0.02, 95% CI = [-0.06, 0.02], *t*(17) = -1.21, *p* = 0.244, *d* = -0.28. B) Relationship between duration judgments and dissimilarity (1 - Event Similarity) for the bilateral hippocampus (left) and right hippocampus (right). Plotted from a model predicting time judgment as a function of: condition, a main effect of a main effect of hippocampal dissimilarity, and a by-participant random effect of condition. Bilateral hippocampus: event dissimilarity significantly predicted duration judgments (more than condition alone): 𝛽 = 0.29; *χ^2^*(1)= 4.67, *p* = 0.031. Right hippocampus: event dissimilarity did not significantly predict duration judgment: 𝛽 = 0.14; *χ^2^*(1)= 1.17, *p* = 0.280. ~p < 0.10; * p < 0.05
